# Estimation of effective population size and effective number of breeders in an abundant and heavily exploited marine teleost

**DOI:** 10.1101/2023.12.17.572092

**Authors:** Andrea Bertram, Justin Bell, Chris Brauer, David Fairclough, Paul Hamer, Jonathan Sandoval-Castillo, Maren Wellenreuther, Luciano B. Beheregaray

**Affiliations:** Molecular Ecology Laboratory, College of Science and Engineering, Flinders University, Bedford Park, SA, Australia; Victorian Fisheries Authority, Queenscliff, Vic, Australia; Aquatic Sciences and Assessment, Department of Primary Industries and Regional Development, Hillarys, WA, Australia; The New Zealand Institute for Plant and Food Research Limited, Nelson, New Zealand; The School of Biological Sciences, University of Auckland, Auckland, New Zealand Corresponding author: Luciano Beheregaray

**Keywords:** fisheries management, fisheries genomics, effective population size, overfishing, genetic diversity

## Abstract

Obtaining reliable estimates of the effective number of breeders *N*(_b_) and generational effective population size (*N*_e_) for fishery-important species is challenging because they are often iteroparous and highly abundant, which can lead to bias and imprecision. However, recent advances in understanding of these parameters, as well as the development of bias correction methods, have improved the capacity to generate reliable estimates. We utilized samples of both single-cohort young of the year and mixed-age adults from two geographically and genetically isolated stocks of the Australasian snapper (*Chrysophrys auratus*) to investigate the feasibility of generating reliable *N*_b_ and *N*_e_ estimates for a fishery species. Snapper is an abundant, iteroparous broadcast spawning teleost that is heavily exploited by recreational and commercial fisheries. Employing neutral genome-wide SNPs and the linkage-disequilibrium method, we determined that the most reliable *N*_b_ and *N*_e_ estimates could be derived by genotyping at least 200 individuals from a single cohort. Although our estimates made from the mixed-age adult samples were generally lower and less precise than those based on a single cohort, they still proved useful for understanding relative differences in genetic effective size between stocks. The correction formulas applied to adjust for biases due to physical linkage of loci and age structure resulted in substantial upwards modifications of our estimates, demonstrating the importance of applying these bias corrections. Our findings provide important guidelines for estimating*N*_b_ and *N*_e_ for iteroparous species with large populations. This work also highlights the utility of samples originally collected for stock structure and stock assessment work for investigating genetic effective size in fishery-important species.

## Introduction

Interest in estimating the genetic effective population size of exploited marine fishes continues to grow as fisheries scientists and managers pay increasing attention to the genetic state of fish stocks (Hare et al. 2011, Ovenden et al. 2015, Marandel et al. 2019). The two genetic effective size parameters, *N*_b_ (effective number of breeders) and*N*_e_ (effective population size), are applicable in a fisheries management context (Hare et al. 2011). The parameter*N*_b_ refers to the number of successful breeders in a single reproductive cycle, which provides important insight into eco-evolutionary processes taking place during reproduction (Waples 1989, Waples and Antao 2014). This parameter is largely shaped by the number and size of families contributing to the sampled cohort, which is influenced by factors like adult density, mate choice, individual variation in fecundity and reproductive success, and habitat quality and quantity (Whiteley et al. 2015). The parameter *N*_e_ represents generational effective population size, which is defined as the size of an idealized population experiencing the same rate of genetic drift or change in genetic diversity per generation as the focal population (Wright 1931). This parameter is thus valuable for determining the effectiveness of selection and population viability, and for developing hatchery-based supportive breeding programs (Charlesworth 2009, Hare et al. 2011). When*N*_e_ is low, increased rates of genetic drift cause genetic variation to erode, the effectiveness of selection to be reduced and deleterious alleles to become fixed, which all cause reductions in fitness, adaptive potential and the probability of population persistence (Luikart et al. 2010, Hare et al. 2011). In exploited species, selective-or over-harvesting and environmental changes can lower*N*_e_ and *N*_b_ due to impacts on demographic parameters like census population size, sex ratios and variance in reproductive success (Luikart et al. 2010, Hare et al. 2011).

Despite the value of effective size parameters in wildlife management, they have proven difficult to estimate accurately and precisely, particularly in abundant and iteroparous (i.e., multiple reproductive cycles over the course of a lifetime) marine species (Hare et al. 2011). This is because a considerable proportion of the census population size needs to be genotyped (>1%; Marandel et al. 2019), and also because age structure (Waples et al. 2014), population genetic structuring (Neel et al. 2013) and physical linkage (Waples et al. 2016) can bias estimates considerably. However, recent developments in understanding these biases, as well as improvements in effective size and confidence interval estimators, mean that our ability to generate accurate and precise estimates has increased considerably. The temporal and single-sample estimators are the most widely used methods for estimating effective size in marine populations (Marandel et al. 2019).However, the temporal method requires genotyping of at least two samples separated by time intervals much larger than the generation time of the focal population (in the case of species with overlapping generations), which is difficult to achieve for long lived and late maturing species like many exploited marine teleosts (Waples 1989). Single-sample estimators such as the linkage disequilibrium (LD*N*_e_) method have therefore become increasingly popular due to their relative practicality (Marandel et al. 2019). The accuracy and precision of the LD*N*_e_ method has also recently been improved through the inclusion of options for screening out rare alleles and the addition of a jackknife confidence interval estimator (Do et al. 2014, Jones et al. 2016). Although employing large numbers of genetic markers (i.e., 1,000s of SNPs) generally increases precision of effective size estimates, over-precision can occur due to the resulting large number of independent comparisons. However, the jackknife confidence interval estimator can be utilized to reduce such over-precision when using large SNP datasets (Jones et al. 2016).

Despite the value of effective size parameters in wildlife management, they have proven difficult to estimate accurately and precisely, particularly in abundant and iteroparous (i.e., multiple reproductive cycles over the course of a lifetime) marine species (Hare et al. 2011). This is because a considerable proportion of the census population size needs to be genotyped (>1%; Marandel et al. 2019), and also because age structure (Waples et al. 2014), population genetic structuring (Neel et al. 2013) and physical linkage (Waples et al. 2016), can bias estimates considerably. However, recent developments in understanding these biases, as well as improvements in effective size and confidence interval estimators, mean that our ability to generate accurate and precise estimates has increased considerably. The temporal and single-sample estimators are the most widely used methods for estimating effective size in marine populations (Marandel et al. 2019).However, the temporal method requires genotyping of at least two samples separated by time intervals much larger than the generation time of the focal population (in the case of species with overlapping generations), which is difficult to achieve for long-lived species, like many exploited marine teleosts (Waples 1989). Single-sample estimators such as the linkage disequilibrium (LD*N*_e_) method (as implemented in NEESTIMATOR; Do et al. 2014) have therefore become increasingly popular due to their relative practicality (Marandel et al. 2019). The accuracy and precision of the LD*N*_e_ method have also been improved through the inclusion of options for screening out rare alleles and the addition of a jackknife confidence interval estimator (Do et al. 2014, Jones et al. 2016). Although employing a large number of genetic markers (i.e., 1,000s of SNPs) generally increases precision of effective size estimates, over-precision can occur due to the resulting large number of independent comparisons. However, the jackknife confidence interval estimator can be utilized to reduce such over-precision when using large SNP datasets (Jones et al. 2016).

Advances in understanding of the magnitude of the biases caused by age structure and physical linkage has led to the development of correction formulas. developed a formula incorporating information on chromosome number to correct for bias in effective size estimates due to physical linkage. Additionally, Waples et al. (2014) produced formulas integrating information on age at maturity and adult lifespan to correct for biases caused by age structure. Besides these bias corrections, it has also been demonstrated that the best approach for obtaining reliable effective size estimates is to employ a sample of individuals from a single cohort (Waples et al. 2014). Using the LD*N*_e_ method, a sample from a single cohort produces an estimate of*N*_b_ relating to the pool of parents that gave rise to the cohort. Generational*N*_e_ can also be calculated from this*N*_b_ estimate using a formula from Waples et al. (2014) and information on age at maturity and adult lifespan. Although capacity to generate reliable effective size estimates requires basic genetic and life-history information, as well as large samples of individuals of known ages, such resources and data are often readily available for fishery important species.

We investigated the potential to generate robust LD*N*_e_-based effective size estimates in a highly abundant iteroparous species, the Australasian snapper (*Chrysophrys auratus*), using genome-wide SNPs and both single-cohort young of the year (YOY) and mixed-age adult samples. Snapper is a long-lived abundant Sparid that inhabits coastal waters of temperate and subtropical Australia as well as northern New Zealand (Gomon et al. 2008). Throughout its range, snapper is a highly important recreational and commercial species, generating significant economic and social benefits (Steven et al. 2021, Jalali et al. 2022, Moore et al. 2023). Because of its abundance and fishery importance, much is known about the biology of snapper and the status of most stocks is assessed regularly (Parsons et al. 2014, Fowler et al. 2021). This means that considerable resources are readily available for estimating the effective size of snapper stocks, including a reference genome, tissue samples from a range of life stages, information on stock structure, as well as population specific data on age at maturity and longevity (Parsons et al. 2014, Catanach et al. 2019, Fowler et al. 2021).

Here we utilized tissue samples taken from snapper belonging to two genetically distinct and geographically isolated stocks in southern Australia that were originally sampled for stock assessment purposes and for population genetic structure work (Conron et al. 2020, Fairclough et al. 2021, Bertram et al. 2022, Bertram et al. 2023). For the two stocks, both single-cohort YOY and adults of mixed ages were available for effective size estimation, allowing us to generate estimates of both *N*_b_ in a single reproductive cycle and generational *N*_e_ using the LD*N*_e_ method. We also compare generational *N*_e_ made from the YOY based *N*_b_ estimates and the samples of mixed-age adults. We apply the bias adjustments to our effective size estimates to account for physical linkage (based on chromosome number; Waples et al. 2016) and age structure (based on age at maturity and adult lifespan; Waples et al. 2014, Waples et al. 2018a). To our knowledge, this is the first study to compare empirical *N*_e_ estimates from both single-cohort and mixed-age adult samples in a highly abundant teleost using genome-wide SNPs. Thus, our study improves understanding of the performance of the LD*N*_e_ method across samples of highly abundant, iteroparous species with different age compositions.

## Materials & Methods

### Sampling

Population genomic work has shown that the Australian locations selected in this study are represented by two geographically and genetically isolated snapper populations, known as the south-west (Bertram et al. 2022) and the south-east (Bertram et al. 2023) stocks. Muscle samples were obtained from 202 0+ aged YOY snapper recruits from the south-west stock (Cockburn Sound), as well as 202 YOY from the south-east stock (Port Phillip Bay; Table 1, Fig. 1). These YOY snapper were originally collected for annual recruitment surveys via trawling by the Department of Primary Industries and Regional Development Western Australia and the Victorian Fisheries Authority. The south-west YOY hatched during the breeding season of 2016, while the south-east recruits hatched during the breeding season of 2017/18. Muscle or fin samples were also obtained from adults of mixed ages belonging to the stocks during 2011, 2014, 2018 and 2019 (Table 1). These adult snapper, which were landed by commercial or recreational fishermen, or by fisheries researchers as part of fisheries independent surveys, were originally sampled for population genetic structure work (Bertram et al. 2022, Gardner et al. 2022, Bertram et al. 2023). The south-west adult sample contained 150 individuals caught in Cockburn Sound, Busselton and Albany (Table 1, Figure 1; Bertram et al. 2022). The south-east adult sample included 185 individuals caught in Kingston SE, Portland, Port Phillip Bay and Western Port Bay (Table 1, Figure 1; Bertram et al. 2023). Where possible, biological data on age and length were obtained for sampled fish (Table 1). Tissue samples were placed in 100% ethanol and stored at −20°C until DNA extraction.

**Figure 1.**
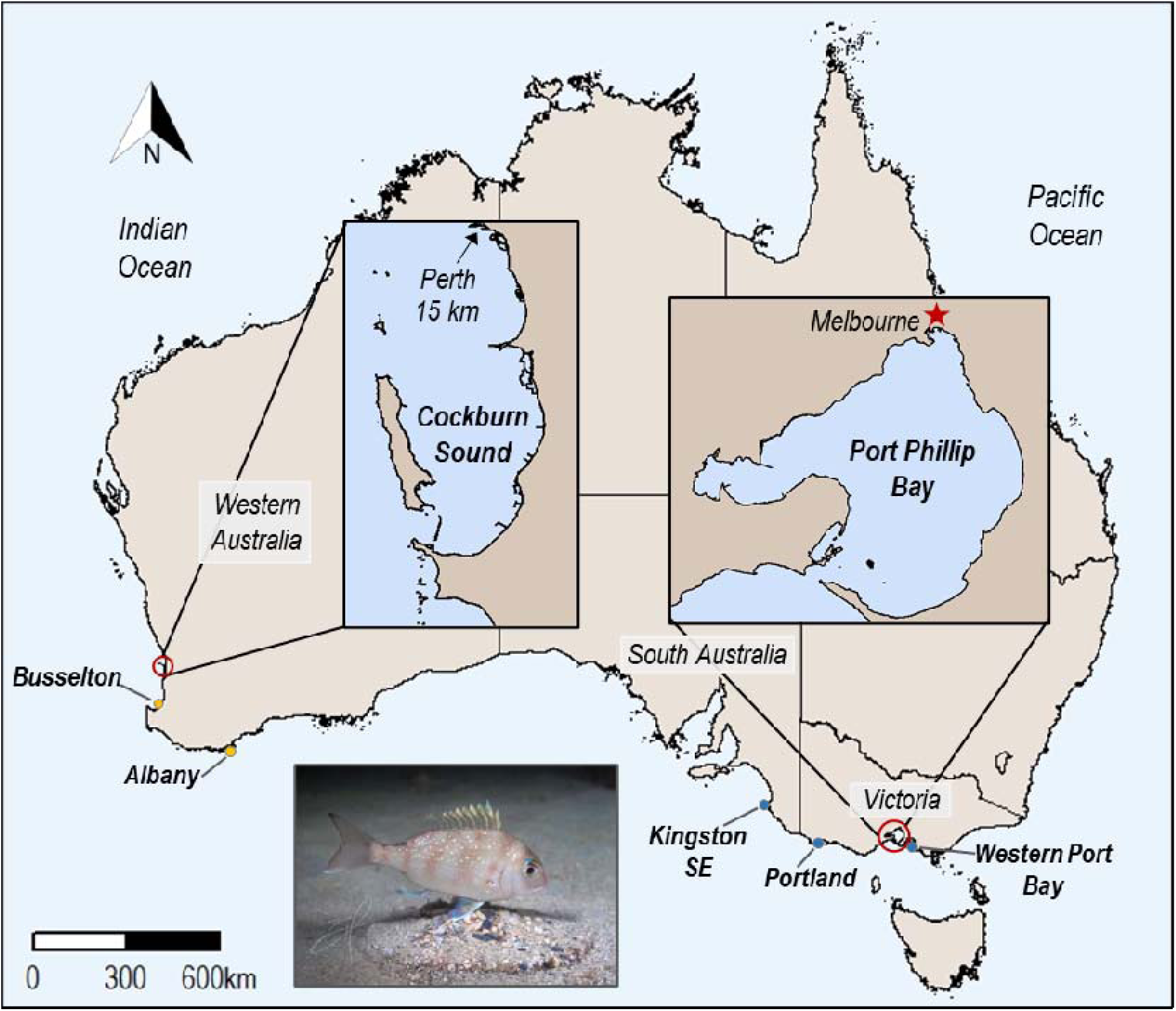
Map of sampling locations for the young of the year (YOY) and adult snapper (*Chrysophrys auratus*) from south-west and south-east stocks. For the south-west, the YOY were obtained from Cockburn Sound (*n* = 202), while the adults were sourced from Cockburn Sound, Busselton, and Albany (*n* = 147). For the south-east, the YOY were obtained from Port Phillip Bay (*n* = 202), while the adults were sourced from Kingston SE, Portland, Port Phillip Bay, and Western Port Bay *n*(= 184). Photo of YOY snapper taken in Cockburn Sound courtesy of Brian Hoehn .

**Table 1.**
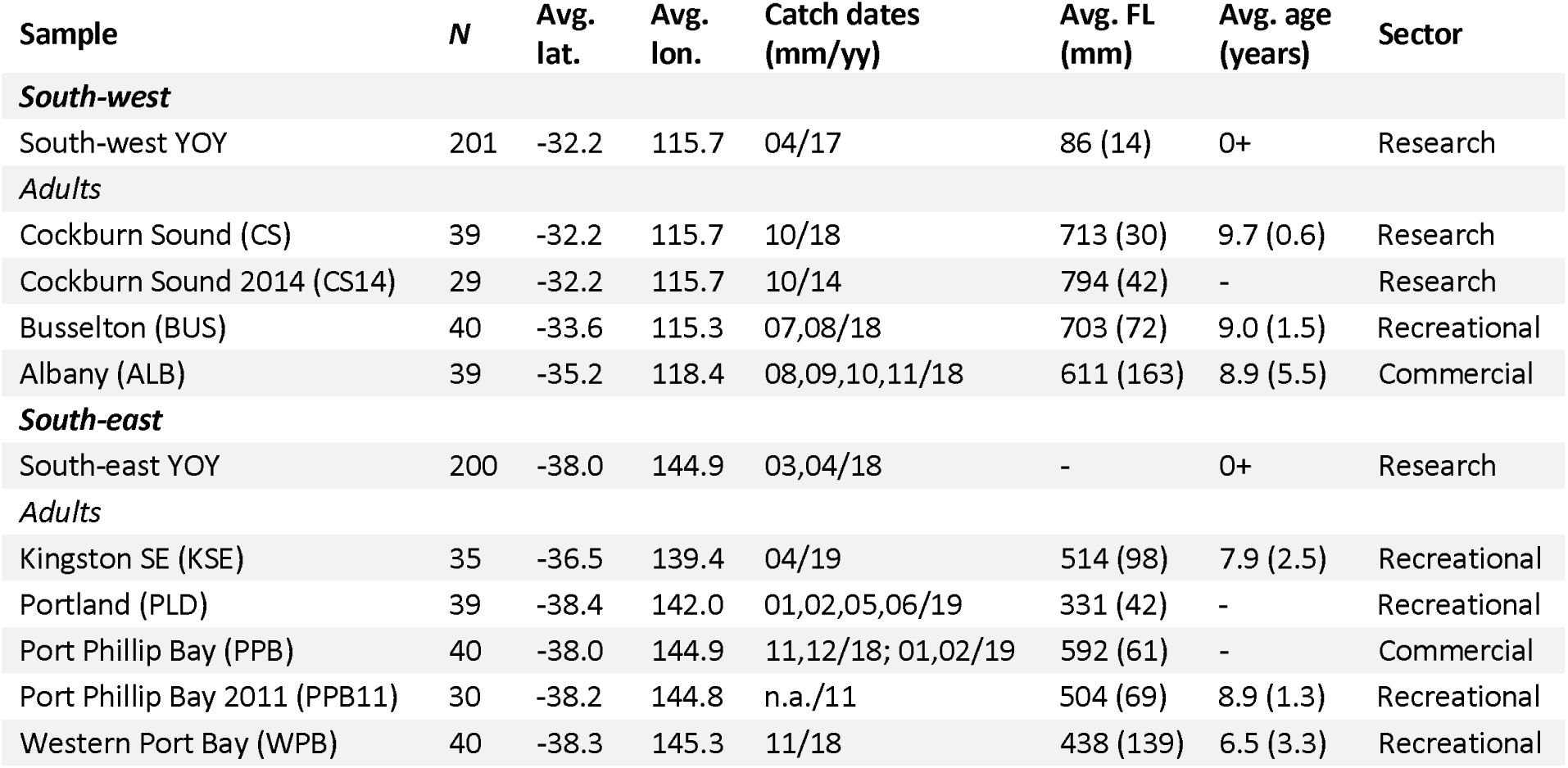
Catch and biological data for the young of the year (YOY) and adult snapper samples from the south-west and south-east snapper stocks in Australia. Average fork lengths (FL) and ages are followed by standard deviations in parentheses. Sample sizes represent the numbers of individuals after removing those with >20% missing data.

### DNA extraction, library preparation and sequencing

For all samples, DNA was extracted using a modified salting-out protocol (Sunnucks and Hales 1996). DNA quality was assessed with agarose gel electrophoresis (1% TBE gel) and extracts were quantified with Qubit v2.0 (Life Technologies). Double-digest restriction site-associated DNA (ddRAD) libraries were then prepared using a protocol modified from Peterson et al. (2012), as detailed in Brauer et al. (2016). Briefly, eight ddRAD libraries, with each comprising 96 DNA samples, were prepared for sequencing. Approximately 200 ng of genomic DNA per sample was digested using the restriction enzymes Sbfl-HF and Msel (New England Biolabs). One of 96 unique 6 bp barcodes was then ligated to each individual sample before being pooled into lots of 12. Using a Pippin Prep (Sage Science), DNA fragments between 300 and 800 bp were selected from each pool. Following PCR of the size selected DNA fragments, size distribution was examined using a 2,100 Bioanalyser (Agilent Technologies) and quantification was carried out with Qubit. Libraries were sequenced on eight lanes of an Illumina HiSeq 4,000 (150 bp paired end) at Novogene (Hong Kong). Replicates were included in each pool of 96 samples for quantification of genotyping and sequencing errors.

### Bioinformatics

Raw sequence reads were demultiplexed using the*process_radtags* module of STACKS 2 (Catchen et al. 2013). Barcodes, restriction sites, and RAD tags were subsequently trimmed from reads with TRIMMOMATIC 0.36 (Bolger et al. 2014). Trimmed sequence reads were mapped to a high-quality snapper genome (Catanach et al. 2019) using BOWTIE 2 (Langmead and Salzberg 2012) before calling single nucleotide polymorphisms (SNPs) with BCFTOOLS 1.16 (Narasimhan et al. 2016). Using VCFTOOLS 0.1.16 (Danecek et al. 2011), individuals with >20% missing data were identified and subsequently excluded before conducting the filtering steps detailed in Table S1. All utilized scripts are available at https://github.com/Yuma248/SNPcallingPipe.

### Categorising neutral loci

Loci under selection are expected to bias effective size estimates (Waples et al. 2016). Therefore, we removed loci from our dataset determined to be under selection based on an analysis using BAYESCAN 2.1 (Foll and Gaggiotti 2008). Twenty pilot runs were undertaken, each with 5,000 iterations, followed by 100,000 iterations with a burn-in length of 50,000 iterations. Outlier loci were identified using a 5% false discovery rate using prior odds of 10.

### Population genomic structure

To confirm that the YOY and adult samples taken from the same geographical region belong to the same genetic population, thereby allowing for effective size estimates to be directly compared, we tested for the presence of population genetic structuring using the maximum-likelihood approach of ADMIXTURE 1.3 (Alexander et al. 2009, Alexander and Lange 2011). We used the software to perform a 5-fold cross-validation for the*K* values 1–5. Ancestry proportions for the most likely*K* value were visualized using GGPLOT 3.3.3 (Wickham 2016) in R.

### Genomic diversity

The genome-wide genetic diversity parameters observed heterozygosity (*H*_O_), expected heterozygosity (*H*_E_), percent polymorphic loci (%PL) and the *F*_IS_ population inbreeding coefficient, were calculated for the YOY and adult samples using the*populations* module in STACKS 2 (Catchen et al. 2013).

### Effective size estimation (N***_b_*** and N***_e_***)

Following Waples et al. (2014), linkage disequilibrium (LD) based effective size estimates based on a single-cohort reflect *N*_b_ in one reproductive cycle, while those based on adults of mixed ages reflect *N*_e_ per generation. Using the LD*N*_e_ method in NEESTIMATOR 2.1 (Do et al. 2014), we estimated *N*_b_ for the south-west and south-east stocks from the YOY samples (which were also subsequently converted to *N*_e_ estimates using equation three detailed below), and estimated*N*_e_ per generation from the two mixed-age adult samples. For the adults,*N*_e_ was calculated both including and excluding the samples collected prior to 2018 to assess the effects of adding extra cohorts on our estimates. We ran the software using the no singleton alleles option and calculated 95% confidence intervals (CIs) using a jackknife method that accounts for pseudoreplication due to linkage and overlapping loci (Jones et al. 2016). Bias corrections were applied to the resulting estimates and their CIs to account for both physical linkage and age structure. The first correction, which was applied to both the raw single-cohort *N*_b_ and the mixed-age *N*_e_ estimates, adjusts for downward bias due to physical linkage, which cannot be fully accounted for by r^2^ filtering methods due to difficulties with identifying loosely linked loci (Waples et al. 2016):

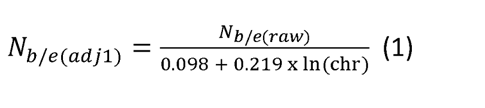

The above formula approximately accounts for the increased LD of linked loci, occurring due to limited recombination, using the number of haploid chromosomes (24 for snapper; Ashton et al. 2019). The process of recombination, including chromosome number, is strongly negatively associated with the magnitude of the bias in *N*_e_ due to linked loci (Waples et al. 2016).

Next, the *N*_b(adj1)_ estimates were adjusted to account for bias due to age structure using information on adult life span (*AL*) and age at maturity (α; Waples et al. 2014):

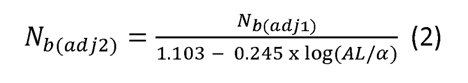

Since the longevity of snapper in Australia is ∼40 years (Norriss and Crisafulli 2010), the AL values used for each stock were 40 minus α estimates. For the south-west YOY sample, α was set to 5.7 (Wakefield et al. 2015) and *AL* to 34.3, while for the south-east YOY sample, an α of 4.9 and an*AL* of 35.1 was used. From these *N*_b(adj2)_ estimates, *N*_e_ per generation was estimated using the same two life history parameters following Waples et al. (2014):

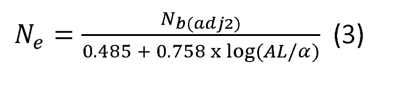

The *N*_e(adj1)_ estimates made from the mixed-age adults were adjusted upwards to account for expected downwards bias due to age structure using the approach of (Waples et al. 2018a) for bluefin tuna. According to Waples et al. (2014),*N*_e_ estimates based on mixed-age samples of iteroparous species with life history traits comparable to snapper (e.g., *AL* and α) are likely downwardly biased by ∼20%. As a result, we adjusted the two*N*_e(adj1)_ estimates upwards by dividing them by 0.8.

## Results

### SNP genotyping

After completing strict quality filtering (Table S1), the YOY and adult datasets comprised 6,839 SNPs. Twenty-one SNP loci determined to be under selection were removed, leaving 6,818 neutral SNPs for our analyses. Of the 753 genotyped snapper, seven were found to have >20% missing data (three YOY and four adults) and therefore were subsequently removed (Table 1). The remaining 401 YOY (201 from the south-west; 200 from the south-east) had an average of 0.6% missing data (range: 0.01–14.84%), while the remaining 331 adults (147 from the south-west; 184 from the south-east) had an average of 0.2% missing data (range: 0–1.5%).

### Population genomic structure

Our ADMIXTURE analysis confirmed a lack of population genomic structure between the YOY and adult samples from the south-west, as well as between the YOY and adult samples from the south-east (*K* = 2 most supported; Figure S2). This indicates that the YOY and adult samples taken from the same geographical region belong to the same genetic populations, allowing for subsequent comparisons of generational *N*_e_ estimates. Additionally, the analysis indicated a lack of temporal genetic structure between the adult samples within each region collected at different time periods (i.e., between the 2014 and 2018 south-west fish, and the 2011 and 2018/19 south-east fish).

### Genomic diversity

Genomic diversity was high and similar across the different snapper samples, with the percentage of polymorphic loci (%PL) ranging between 99.5% (south-west and south-east adults) and 99.7% (south-east YOY), and observed and expected heterozygosity (*H*_O_ and *H*_E_) ranging between 0.206 (south-west YOY) and 0.208 (south-east YOY and adults), and 0.209 (south-west YOY and adults) and 0.210 (south-east YOY and adults), respectively (Table 2). Values of*F*_IS_ were close to zero for all samples, ranging between 0.012 (south-east adults) and 0.020 (south-west YOY; Table 2). The south-east samples had slightly higher *H*_O_ and *H*_E_ and slightly lower *F*_IS_ than the south-west samples. The diversity statistics quoted above for the adult samples were generated without the fish collected prior to 2018. However, genomic diversity was highly similar between the samples excluding and including these additional fish (Table 2).

**Table 2.** Summary of sample sizes and genomic diversity (based on 6,818 SNPs) for the young of the year (YOY) and adult snapper samples excluding (adults 1) and including (adults 2) the extra fish obtained prior to 2018. Abbreviations are percent polymorphic loci, %PL; expected heterozygosity, *H*_E_; observed heterozygosity, *H*_O_; inbreeding coefficient, *F*_IS_.

| Sample | *N* | %PL | $H_O$ | $H_E$ | $F_{IS}$ |
| --- | --- | --- | --- | --- | --- |
| South-west YOY | 201 | 99.6 | 0.206 | 0.209 | 0.020 |
| South-east YOY | 200 | 99.7 | 0.208 | 0.210 | 0.017 |
| South-west adults 1 | 118 | 99.5 | 0.207 | 0.209 | 0.018 |
| South-west adults 2 | 147 | 99.6 | 0.207 | 0.209 | 0.020 |
| South-east adults 1 | 154 | 99.5 | 0.208 | 0.210 | 0.012 |
| South-east adults 2 | 184 | 99.6 | 0.209 | 0.210 | 0.013 |

### Estimates of N***_b_*** in a single reproductive cycle

The bias corrections to account for physical linkage and age structure resulted in upwards adjustments of the raw south-west and south-east YOY*N*_b_ estimates and their 95% confidence intervals (CIs) by 41% and 43%, respectively (Table 3). Adjusted*N*_b_ was greater for the south-west than the south-east YOY sample (3,754 vs 2,684). The lower bounds of the 95% CIs for the south-west and south-east YOY *N*_b_ estimates were similar (1,702 vs 1,362), while their upper bounds were indeterminate (i.e., infinite) and 35,587, respectively.

**Table 3.** Raw and adjusted effective size estimates based on 6,818 SNPs for the two young of the year (YOY) snapper samples. Raw estimates were calculated using the linkage disequilibrium method in NEESTIMATOR 2.1 (Waples and Do 2008, Do et al. 2014), and reflect the effective number of breeders (*N*_b_) in one reproductive cycle. The first and second adjustments made to the *N*_b_ estimates were applied to account for bias due to physical linkage of loci and age structure, respectively. Generational effective population size (*N*_e_) was calculated from the final adjusted*N*_b_ estimates using an equation from Waples et al. (2014) that incorporates information on two life history traits (adult lifespan and age at maturity). In parentheses are 95% confidence intervals generated using the jackknife method in NEESTIMATOR.

| Sample | $N$ | $N_{b(\text{raw})}$ | $N_{b(\text{adj}1)}$ | $N_{b(\text{adj}2)}$ | $N_e$ |
| --- | --- | --- | --- | --- | --- |
| South-west YOY | 201 | 2,670<br>(1,210–inf) | 3,363<br>(1,524–inf) | 3,754<br>(1,702–inf) | 3,333<br>(1,511–inf) |
| South-east YOY | 200 | 1,875<br>(951–24,854) | 2,361<br>(1,198–31,302) | 2,684<br>(1,362–35,587) | 2,282<br>(1,158–30,256) |

### Estimates of generational N***_e_***

Generational *N*_e_ based on the south-west and south-east YOY samples, which we calculated from the adjusted *N*_b_ estimates, was 3,333 and 2,282, respectively (95% CIs: 1,511–inf and 1,158– 30,256; Table 3, Fig. 2). The bias adjustments (to account for physical linkage and age structure) applied to the raw *N*_e_ estimates for the south-west and south-east adult samples resulted in upwards modifications of 74% (Table 4). Adjusted*N*_e_ was similar across the adult samples, albeit slightly higher for the south-east than the south-west (1,958 vs 1,834; Table 3, Fig. 2). Like the YOY based *N*_e_ estimates, the lower bounds of the 95% CIs for the adult*N*_e_ estimates were similar (south-west, 779; south-east, 953). Additionally, the 95% CIs for south-west adult*N*_e_ estimate also had an indeterminate upper limit (i.e., infinite). The upper limit of the 95% CIs for the south-east adult *N*_e_ estimate was 126,951. *N*_e_ estimates including the fish collected prior to 2018 were slightly higher for both stocks (south-west, 2,631; south-east, 2,687; Table 3, Fig. 2). The lower bounds of the 95% CIs for these estimates were also higher (south-west, 1,183; south-east, 1,339). The south-west estimate continued to exhibit an indeterminate upper limit, while the south-east upper limit was less than half of that obtained for the smaller sample (54,282).

**Table 4.** Raw and adjusted effective size estimates based on 6,818 SNPs for the mixed-age adult snapper samples excluding (adults 1) and including (adults 2) the extra fish obtained prior to 2018. Raw estimates were calculated using the linkage disequilibrium method in NEESTIMATOR 2.1 (Waples and Do 2008, Do et al. 2014) and reflect generational effective population size *N*(_e_). The first and second adjustments applied to the*N*_e_ estimates were to account for bias due to physical linkage of loci and age structure, respectively. In parentheses are 95% confidence intervals generated using the jackknife method in NEESTIMATOR.

| Sample | *N* | $N_{e(\text{raw})}$ | $N_{e(\text{adj1})}$ | $N_{e(\text{adj2})}$ |
| --- | --- | --- | --- | --- |
| South-west adults 1 | 118 | 988<br>(444 - inf) | 1,244<br>(559 - inf) | 1,716<br>(771 - inf) |
| South-west adults 2 | 147 | 1,515 (681 - inf) | 1,907 (858 - inf) | 2,631<br>(1,183 - inf) |
| South-east adults 1 | 154 | 1,129<br>(571 - 13,636) | 1,422<br>(719 - 17,174) | 1,962<br>(992 - 23,689) |
| South-east adults 2 | 184 | 1,547<br>(771 - 31,247) | 1,948<br>(971 - 39,354) | 2,687<br>(1,339 - 54,282) |

**Figure 2.**
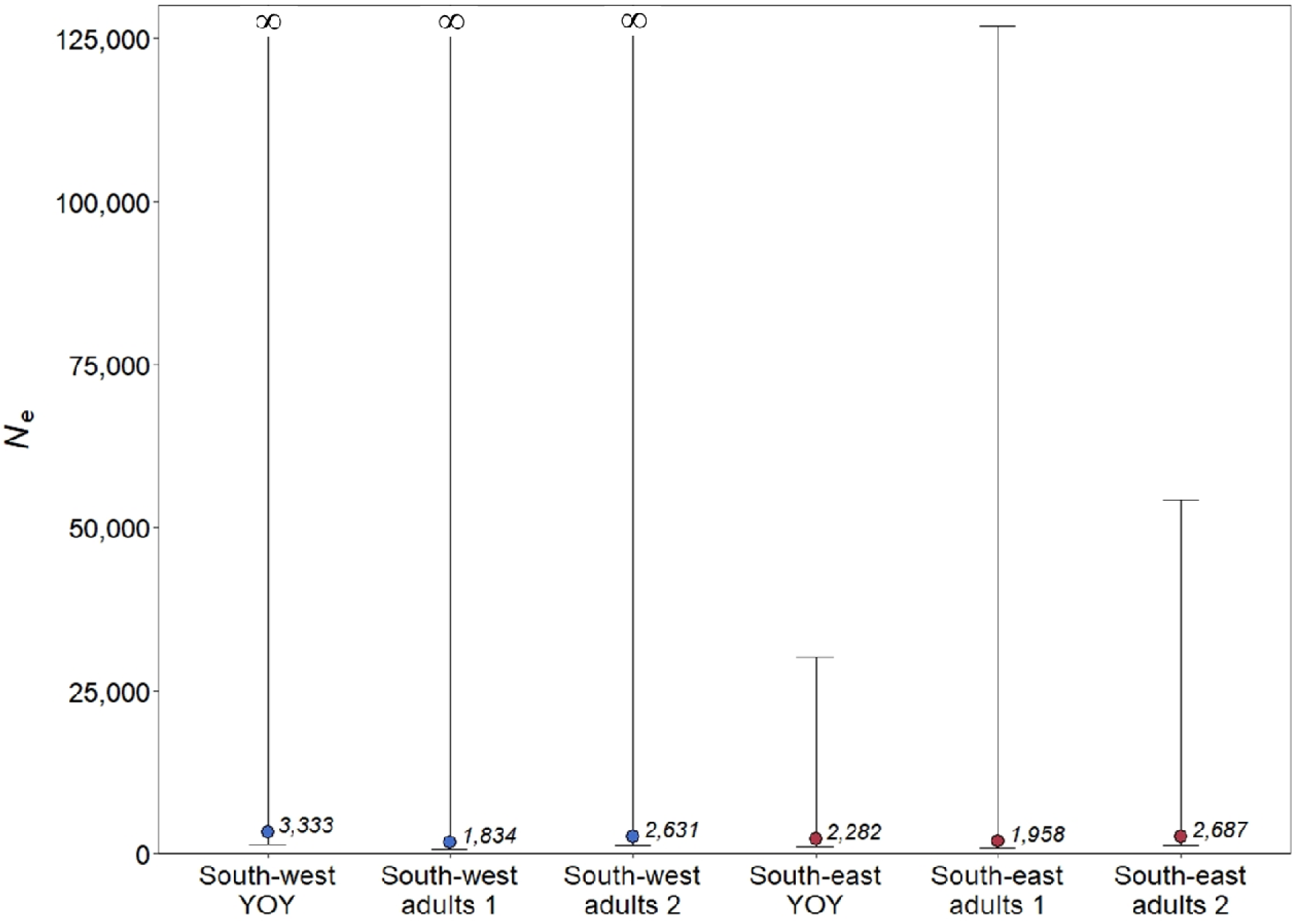
Comparison of generational effective population size (*N*_e_) based on 6,818 SNPs for the young of the year (YOY) and adult snapper from the south-west (blue) and south-east (red) stocks in Australia. The first and second estimates for the mixed-age adult samples exclude and include the extra fish obtained prior to 2018 respectively. Data labels represent the*N*_e_ estimates after correcting for biases due to physical linkage and age structure.

## Discussion

There is significant value in estimating the genetic effective size (*N*_e_ and *N*_b_) of wildlife populations for conservation and management purposes. Although estimating*N*_e_ and *N*_b_ is challenging, particularly in abundant and iteroparous species, advances in our understanding of these parameters have improved our ability to generate reliable effective size estimates. We assessed the potential to generate reliable *N*_e_ and *N*_b_ estimates for an abundant, iteroparous species using genome-wide SNPs and samples from both single-cohort young of the year (YOY) and adults of mixed ages. To explore this, we focused on two genetically distinct Australian stocks of the fishery-important teleost, the Australasian snapper (*Chrysophrys auratus*), utilizing fish originally sampled for stock assessment and population genetic structure work. Linkage disequilibrium (LD)*N*_e_ estimates based on single-cohort YOY were generally higher and more precise than those based on the adults of mixed ages. Upper bounds of the 95% confidence intervals around our*N*_e_ and *N*_b_ estimates were achieved for one of the two snapper stocks. Our study indicates that it is possible to generate reliable genetic effective size estimates for abundant, iteroparous species using large samples and genome-wide SNPs, especially if samples from a single cohort are available for genotyping and if the relevant bias corrections are applied. This work also demonstrates the potential additional uses of specimens originally collected to address other research questions. The design and results of this study can inform the development of strategies for obtaining reliable genetic effective size estimates for other abundant, iteroparous species.

### Genetic effective size bias corrections

The corrections applied to account for bias due to physical linkage of loci and age structure resulted in considerable upwards modifications to our effective size estimates. Estimates based on the south-west and south-east YOY samples were adjusted upwards by 41% and 43% respectively, while those based on the mixed-age adult samples were adjusted upwards by 74%. Species with ∼60 chromosomes will produce genetic effective size estimates with minimal bias due to physical linkage of loci (Waples et al. 2016). However, downward bias occurs in species like snapper that have <60 chromosomes, particularly if chromosomes are short in length (<100 cM), because smaller genomes contain fewer independently assorting loci. With respect to age structure, downward bias in effective size estimates increases as the ratio between adult lifespan and age at maturity increases (Waples et al. 2014). Therefore, effective size estimates can be highly inaccurate if these bias corrections are not applied, particularly in species that have less than or more than sixty chromosomes, and in species that have long life spans with early maturity.

### Effective number of breeders (N***_b_***) in a single reproductive cycle

The most straightforward approach for obtaining reliable genetic effective size estimates with the LD*N*_e_ method is to genotype a large sample of a single cohort (Waples et al. 2014). The resulting estimates reflect the effective number of breeders (*N*_b_) that gave rise to the sampled cohort. Our *N*_b_ estimates based on the single-cohort YOY snapper from the south-west and south-east stocks were 3,754 and 2,684, respectively (95% CIs: 1,702–inf and 1,362–35,587). These results suggest that a greater number of individuals successfully contributed to the 2016 south-west cohort than the 2017/18 south-east cohort. This is the first study to generate empirical estimates of*N*_b_ for snapper. Factors that can influence the number of individuals that successfully contribute to a cohort include individual variation in fecundity, population density, sexual selection, spawning and nursery habitat quality and quantity, and the suitability of environmental conditions for spawning and larval survival (Whiteley et al. 2015). Long-term studies on*N*_b_ may be valuable for understanding the factors influencing interannual variation in individual reproductive contribution and therefore may facilitate predicting the impacts of anthropogenic development, different harvest strategies and environmental changes on stock resilience and productivity (Bacles et al. 2018, Luikart et al. 2021). For example, a long-term study by Whiteley et al. (2015) determined that interannual changes in *N*_b_ in two brook trout populations were significantly correlated with temporal variation in stream flow.*N*_b_ monitoring may be particularly feasible in situations where regular sampling of recruits is carried out as part of an existing stock or population monitoring strategy, as is the case for the snapper stocks in this study.

### Generational effective population size (N***_e_***)

Generational effective population size (*N*_e_) refers to the theoretical size of a population facing the same rate of genetic drift or change in genetic diversity per generation as the one in question (Wright 1931). This parameter is of the most informative in wildlife conservation and management as it indicates the vulnerability of a population to environmental changes and exploitation. We calculated *N*_e_ for the two Australian snapper populations using two approaches. First, we calculated *N*_e_ from our YOY based *N*_b_ estimates using population specific information on adult lifespan and age at maturity (Waples et al. 2014). Second, we estimated*N*_e_ from samples of mixed-age adults and used the ratios of observed to expected *N*_e_ for species similar to snapper from Waples et al. (2014) to roughly correct for bias due to age structure. Our*N*_e_ estimates based on the YOY samples were higher and more precise than those based on the mixed-age adult samples. The YOY based *N*_e_ estimates were 3,333 and 2,282 for the south-west and south-east snapper stocks, respectively (95% CIs 1,511–inf and 1,158–30,256), while those based on the south-west and south-east mixed-age adult samples (sampled in 2018 and 2019) were 1,958 and 1,834, respectively (95% CIs: 779–inf and 953–126,951). Including the additional adult snapper collected for prior population genetic work (Cockburn Sound in 2014, *n* = 29; Port Phillip Bay in 2011, *n* = 30), produced *N*_e_ estimates for both stocks that were higher (south-west, 2,631; south-east, 2,687) and more precise (95% CIs 1,183–inf; 1,339–54,282), likely due to the increases in sample sizes. The inclusion of additional cohorts therefore did not result in extra downward bias due to age structure. These *N*_e_ estimates are perhaps more reliable than those generated without these additional samples, as they exhibited precision more comparable to the estimates based on the YOY samples, demonstrating the importance of sample size in generating reliable*N*_e_ estimates.

Similarities between the *N*_e_ estimates based on the different sample types increase our confidence in their validity. Our *N*_e_ estimates for the south-west stock were generally higher than those obtained for the south-east stock, and the upper 95% CI for the south-west estimates were both indeterminate (i.e., infinite). Since the south-west point estimates were generally highest and precision is inversely related to true*N*_e_ (Waples and Do 2010), we can conclude that the*N*_e_ of the south-west stock, when sampled, was larger than that of the south-east stock. This does not necessarily mean that the former stock has a larger census population size than the latter stock, as no simple relationship between *N*_e_ and census population size has been determined and the ratio between the two parameters can vary between populations of the same species (Palstra and Fraser 2012, Pierson et al. 2018). In fact, the relative biomass of the south-west stock is considered to be lower than that of the south-east stock (depleted vs. sustainable, respectively; Fowler et al. 2021), and historic landings have generally been higher in the latter (Conron et al. 2020, Fairclough et al. 2021). Additionally, in 2018, the abundance of YOY in Port Phillip Bay (the primary nursery area for the south-east stock) was the highest since recruitment surveys began 30 years ago (Bell et al. 2021). Although no spawning biomass estimates are available for the south-west snapper stock, population dynamic modelling estimated that the spawning biomass of the south-east stock (from 2016) was >1,000,000 spawning individuals (Hamer et al. 2019). This suggests that the*N*_e_ and *N*_b_ of the south-east may be a small proportion of the census population size, and therefore that individual variation in reproductive contribution is likely to be substantial. This hypothesis is consistent with the vast interannual variation in spawning success of snapper in Port Phillip Bay, where larval survival and juvenile recruitment is linked to changes in abundance and composition of their planktonic diet (Murphy et al. 2013).

Since snapper appear to have colonized the Australian coastline in an east to west direction, the south-west and its two adjacent stocks are likely the most recently formed (Bertram et al. unpublished). These three stocks (the mid-west, south-west and south-coast stocks) are therefore weakly differentiated, and they are also not completely contemporarily isolated. As a result, it is possible that the south-west estimates reflect the effective size of a broader region, or are inflated due to contemporary gene flow from these weakly differentiated adjacent stocks (Waples and England 2011, Neel et al. 2013). Alternatively, the lower*N*_e_ of the south-east compared to the south-west could be due to higher individual variance in reproductive success in the former stock.

Our results suggest that while samples of mixed-age adults are valuable for assessing relative population differences in *N*_e_, they can produce more downwardly biased and less precise estimates than samples from a single cohort. Waples et al. (2014) showed that downward bias of*N*_e_ estimates made from adults of mixed ages increases as the ratio between adult lifespan and generation length increases. Therefore, if the study species is long lived, matures early and reproduces for its entire mature lifespan (like snapper and many other marine fishes), then considerable downward bias in *N*_e_ estimates is expected if they are generated from adults of mixed ages. Compared with a single cohort, *N*_e_ estimates made from mixed-age adults are more difficult to adjust accurately to account for bias due to age structure. Therefore, as already recommended by others, we advise that the most suitable approach for estimating*N*_e_ in species like snapper is to base calculations on a single cohort. Based on the guidelines of Frankham et al. (2014), our single-cohort based *N*_e_ estimates for the south-west and south-east snapper stocks are likely sufficiently large to avoid inbreeding (*N*_e_ >100) and loss of adaptive potential (*N*_e_ >1,000).

### Effects of population genetic structure on effective size estimates

Downward bias in *N*_e_ is expected when population genetic structure occurs within the genotyped sample. This is because the inclusion of genetically divergent individuals generally results in downwardly biased *N*_e_ estimates due to mixture LD (Neel et al. 2013). It is possible that the weak signal of isolation by distance in the south-west (Bertram et al. 2022) caused additional downward bias of our mixed-age adult *N*_e_ estimates. However, we believe the impact of this slight genetic structuring on our *N*_e_ estimates was low, since the most differentiated south-west samples had an *F*_ST_ of only 0.003 (Cockburn Sound vs. Albany; Bertram et al. 2022). Additionally,*F*_IS_ was no higher for the adult sample than for the YOY sample that was obtained from one location (i.e., Cockburn Sound). We would expect inflated*F*_IS_ if genetic structuring due to the Wahlund effect was significant enough to downwardly bias our *N*_e_ estimate (Neel et al. 2013, Waples et al. 2018b). Alternatively, the lower *N*_e_ estimate obtained from the mixed-age adult sample could be due to its age composition causing more downward bias than expected or due to the slightly different time period the estimate relates to (Waples 2005).

### Comparisons with effective size estimates of other species and previous snapper studies

Our *N*_e_ estimates are within the range of those reported for marine species exhibiting very large populations (Marandel et al. 2019), and are similar to*N*_e_ estimates generated for species with reproductive strategies comparable to snapper, including the giant black tiger shrimp*(Penaeus monodon*; Vu et al. 2020), Sydney rock oyster (*Saccostrea glomerata*; O’Hare et al. 2021), green abalone (*Haliotis fulgens*; Gruenthal et al. 2014), Pacific cod (*Gadus macrocephalus*; Suda et al. 2017), white hake (*Urophycis tenuis*; Roy et al. 2012) and redbelly yellowtail fusilier (*Caesio cuning*; Ackiss et al. 2018). As expected, the two snapper *N*_e_ estimates are generally larger than those for anadromous fishes Our *N*_e_ estimates are within the range of those reported for marine species exhibiting very large populations (Marandel et al. 2019), and are similar to*N*_e_ estimates generated for species with reproductive strategies comparable to snapper, including the giant black tiger shrimp (*Penaeus monodon*; Vu et al. 2020), Sydney rock oyster (*Saccostrea glomerata*; O’Hare et al. 2021), green abalone (*Haliotis fulgens*; Gruenthal et al. 2014), Pacific cod (Gadus macrocephalus; Suda et al. 2017), white hake (*Urophycis tenuis*; Roy et al. 2012) and redbelly yellowtail fusilier (*Caesio cuning*; Ackiss et al. 2018). As expected, the two snapper *N*_e_ estimates are generally larger than those for anadromous fishes (Ferchaud et al. 2016, Barría et al. 2019, Waldman et al. 2019, Miller et al. 2022) and elasmobranchs (Dudgeon and Ovenden 2015, Pazmiño et al. 2017, Reid-Anderson et al. 2019, Venables et al. 2021), and are smaller than those for southern bluefin tuna (*Thunnus maccoyii*; Waples et al. 2018a), albacore tuna and New Zealand hoki (*Macruronus novaezelanidae*; Koot et al. 2021), which support far more productive fisheries than snapper. Although fewer studies have explored*N*_b_ in marine species, due to the close relationship between *N*_e_ and *N*_b_, similar trends to those described above occur between our results and similar studies with regard to in *N*_b_ (Whiteley et al. 2015, Puritz et al. 2016, Waples et al. 2018a, Davenport et al. 2021, King et al. 2023).

*N*_e_ has previously been estimated for snapper in eastern Australia and New Zealand. Morgan et al. (2018) estimated *N*_e_ for samples of mixed-aged adult snapper from nine locations in eastern Australia. However, sample sizes were all <60 individuals and only nine microsatellite DNA markers were used, so three samples produced indeterminate point estimates and all estimates had indeterminate upper 95% CIs. estimated *N*_e_ for adult snapper from Tasman Bay and Hauraki Gulf (*n* = 234 for each site) in New Zealand using seven microsatellite DNA markers. Estimated*N*_e_ was 104 (95% CIs 80–720) for Tasman Bay and 1,164 (95% CIs 157–inf) for Hauraki Gulf. Hauser et al. (2002) suggested that the very low *N*_e_ for Tasman Bay could partly be due to the population being located at the southern edge of snapper’s distribution, which may result in recruitment failure in some years. used nine microsatellite DNA markers to estimate*N*_e_ for snapper within a marine reserve in the Hauraki Gulf. Estimated*N*_e_, which was based on 1,044 mixed-age adults, was 10,488 (95% CIs 2,818–inf). *N*_e_ simulations conducted with NEOGEN software (Blower et al. 2019) suggested that at least 1,500 individuals (∼5% of the marine reserve adult population) would need to be genotyped to generate an estimate with a finite upper 95% CI. As with our results, the above studies indicate that large numbers of individuals need to be genotyped to produce precise*N*_e_ estimates for very large populations.

## Conclusions

Generating reliable genetic effective size estimates for fishery-important species is challenging since they often have large, connected populations with overlapping generations. However, abundant resources are often available for investigating genetic effective size in fishery-important species because they are commonly sampled for stock structure work and for routine stock assessment. Our results indicate that it is feasible to obtain reliable effective size estimates for fishery-important species, particularly if large samples from a single cohort are available for genotyping. However, even if such samples are available, estimates can be inaccurate if adjustments are not made to account for factors like physical linkage of loci and age structure. Our study can be used as a guide for others to generate reliable genetic effective size estimates for abundant, iteroparous species.

## Data Availability Statement

The SNP dataset is available on figshare: XXX to be included after manuscript acceptance.

## Acknowledgements

This work was funded by the Australian Research Council (LP180100756 to LBB), Flinders University and by the PhD scholarships awarded to AB from AJ and IM Naylon and the Playford Trust. We thank all who assisted with sample collection, which included research staff at the Department of Primary Industries and Regional Development Western Australia and Victorian Fisheries Authority, as well as Michelle Gardner (Murdoch University) and members of Portland Sport Fishing Club and the organisers of the 2019 Kingston Offshore Fishing Competition. We are also grateful to Andrea Barceló and Diana-Elena Vornicu at the Molecular Ecology Lab at Flinders University for their assistance in the laboratory.

