## Supplemental File for "Estimation of effective population size and effective number of breeders in an abundant and heavily exploited marine teleost"

**Table S1** Numbers of SNPs retained after each bioinformatic filtering step.

| Step | SNP count |
| --- | --- |
| Raw SNP catalogue | 7,338,377 |
| 80% of individuals, biallelic, >0.03 minor allele frequency | 45,715 |
| Remove indels | 41,513 |
| Read quality (ratio quality/coverage depth >0.2) | 40,935 |
| Mapping quality (>30) | 37,041 |
| High coverage loci (≤mean depth + (2*standard deviation)) | 35,992 |
| Hardy-Weinberg equilibrium in >67% of locations | 31,275 |
| Call error rate (0.95) | 27,794 |
| Linkage disequilibrium (500) | 6,839 |
| Putatively neutral loci | 6,818 |
